# Scaling Variant-Aware Multiplex Primer Design

**DOI:** 10.64898/2026.02.03.703607

**Authors:** Yunheng Han, Christina Boucher

**Affiliations:** Department of Computer and Information Science and Engineering, Herbert Wertheim College of Engineering, University of Florida, 1889 Museum Road, Gainesville, 32611, Florida, USA

**Keywords:** Multiplex PCR, Variant-Aware Primer Design, Primer-Dimer Minimization, Convex Optimization, Combinatorial Optimization

## Abstract

**Motivation:** Robust primer design is essential for reliable multiplex PCR in diverse and evolving pathogen, microbial, and host genomes. Traditional methods optimized for a single reference often fail on emerging variants, leading to reduced efficiency. Variant-aware design seeks primers that remain effective across diverse targets, but this introduces two key challenges: identifying robust candidates and selecting an optimal subset of primers. Although there are methods for the first challenge, namely the Primer Design Region (PDR) optimization problem, existing approaches lack optimality guarantees.

**Results:** We introduce a near-linear algorithm with provable guarantees for efficient PDR optimization. Complementing this, we introduce a reference-free risk model based on Gini impurity that provides a stable, biologically interpretable measure of site-specific variation and yields PDRs that are robust to sequence diversity across datasets without ad hoc smoothing. For the second challenge related to thermodynamic stability, we optimize predicted Δ*G* and cast subset selection as a *k*-partite maximum-weight clique problem (NP-hard). We then design an efficient local-search heuristic with linear time updates. Together, these advances yield a principled, scalable framework for variant-aware primer design. Across Foot-and-Mouth Disease virus and Zika virus datasets, Δ-PRO produces more compact and robust PDR sets and multiplex panels with reduced predicted dimerization compared to existing tools, demonstrating the practical gains of principled and scalable variant-aware primer design for high-throughput multiplex PCR assays.

**Availability:** The proposed methods are implemented in a software package. Our implementation and results are publicly available at https://github.com/yhhan19/variant-aware-primer-design.

**Supplementary information:** Supplementary materials are available online.

## Introduction

Primer sequences are the design variables that set the signal-to-noise ratio for PCR and downstream sequencing Xie et al. [2022]. When primers are suboptimal, off-target amplification increases, yield drops, and read distributions become biases Goodwin et al. [2016], Lear et al. [2018]. These risks escalate in multiplex PCR, where dozens to hundreds of primers must operate in the same reaction, and therefore, must be co-optimized to minimize cross-hybridization. The dominant failure mode is primer-primer binding (also called dimerization), which occurs when complementary bases of primers form duplexes that polymerase can extend, diverting reagents from true targets and producing artifacts that complicate inference Elnifro et al. [2000], Khodakov et al. [2016].

Given the importance of primer design for multiplex PCR, a number of optimization methods and algorithms have been created to select primers in a manner that avoids dimerization. Primer3 Untergasser et al. [2012] and Primer-BLAST Ye et al. [2012] are some early methods that remain reliable tools for selecting individual primers using criteria such as melting temperature, GC content, self-complementarity, and alignment-based specificity. However, their scope is largely limited to single-plex, i.e., they evaluate primers one pair at a time and do not model how a larger set will behave together (e.g., cross-hybridization or dimer formation among primers in a multiplex reaction). Consequently, they can yield high-quality primers in isolation without ensuring that the full panel operates coherently in a shared mix. Some early methods were developed specifically for multiplex PCR and began to explicitly model interactions within primer pools, giving rise to tools tailored for multiplex use cases. These methods include MultiPLX Kaplinski et al. [2005], MuPlex Rachlin et al. [2005], MPprimer Shen et al. [2010], PrimerStation Yamada et al. [2006] and MPD Wingo et al. [2017]. However, many of these methods are no longer maintained and those that are often impose practical constraints, such as reliance on user-specified targets and strict limits on the number of genomes that can be multiplexed at once.

Several recent methods for multiplex PCR have been developed specifically to support many more genomes in a single experiment, but achieving this scale requires more aggressive and systematic optimization to control primer–primer dimerization. PMPrimer Yang et al. [2023] uses multiple sequence alignment with entropy filters to select conserved regions, identifies primers for each of these regions, and then pools all primers. Although PMPrimer was one of the first methods to design primers in a manner that accounts for the conserved loci, it leads to a large number of primers because there is no attempt to optimize the final pooled set. PrimalScheme Kent et al. [2024] provides an accessible pipeline for viral sequencing, but its heuristics can falter on highly diverse templates or when scaling to large target sets. SADDLE Xie et al. [2022] and Olivar Wang et al. [2024] provide a more principled approach to optimize dimerization for multiplex PCR. SADDLE Xie et al. [2022] introduced the concept of a *Badness score* for a set of primers, which aims to quantify dimer risk by combining sequence complementarity, thermodynamic stability, and 3’-end proximity. This Badness score is an alternative to thermodynamic stability Δ*G*, which is traditionally used to measure dimerization. Olivar Wang et al. [2024] also adopts this metric and uses heuristics to minimize aggregate Badness across a set of primers. One of the shortcomings of SADDLE is that variation across the genome is not taken into account when designing the primers. This is significant since primers frequently fail on divergent variants, leading to allele dropout and uneven coverage.

To overcome some of the constraints imposed by current laboratory primer design methods, multiplex primer design must explicitly account for genomic variation. To our knowledge, Olivar Wang et al. [2024] is the first and only existing method that explicitly performs variant-aware multiplex primer design. It constructs a nucleotide-level “risk landscape” by combining local metrics such as the single nucleotide polymorphism (SNP) frequency, GC content, sequence complexity, and non-specificity, then identifies contiguous Primer Design Regions (PDRs) from which robust primers can be drawn. To be clear, the PDRs are defined in order to minimize the “risk” imposed for loci with high heterogeneity. Nevertheless, Olivar’s risk formulation is appealing because it flexibly integrates diverse site-based metrics into a unified *signal vector* of risks, which enables efficient optimization over genomic regions for primer design. However, Olivar uses SNP frequencies or Shannon entropy as underlying variation metrics, both of which have notable limitations. SNP frequency depends on a chosen reference and ignores the full nucleotide distribution, making it unable to distinguish between sites with different frequency distributions. In contrast, Shannon entropy is reference-free but lacks a clear biological interpretation, depends on the logarithm base, and is overly sensitive to rare frequency values. In practice, additional transformations, such as taking the square root, are used to amplify weak signals or smooth variation, but these ad hoc adjustments lack theoretical grounding and require manual tuning for stability across datasets. Furthermore, Olivar’s PDR optimization procedure remains stochastic and heuristic, lacking guarantees on optimality or stability.

To overcome these limitations, we present a variant-aware, multiplex primer design algorithm that builds a nucleotide-level risk landscape and co-optimizes PDRs and primer sets at scale. We refer to our method as Δ-PRO (Δ-guided PRimer Optimization). The contributions of our work are threefold. First, we introduce a biologically interpretable reference-free variation metric based on Gini impurity, providing a stable probabilistic estimate of site diversity. The experimental results show that the Gini-based variation metric achieves greater robustness, while remaining stable across datasets, unlike entropy or SNP-based metrics that require additional smoothing transformations, whose performance can fluctuate. Next, we propose a dynamic programming algorithm that optimizes the PDR, providing provable global optimality and near-linear efficiency. Finally, we directly minimize dimerization based on predicted thermodynamic stability (Δ*G*) through a graph-based formulation, where primers are represented as vertices in a *k*-partite graph and edges are weighted by their Δ*G* value. This corresponds to a maximum-weight clique problem, which is NP-hard, and we address it using efficient local search heuristics that explore the entire 𝒪(*p*) neighborhood in 𝒪(*p*) time, where *p* denotes the number of candidate primers, enabling efficient exploration of the search space and scalable optimization for large multiplex panels. We demonstrate the practical utility of our method through experimental studies on the Foot-and-Mouth Disease virus and Zika virus datasets, where we compare its performance with that of existing approaches. Our implementation and results are publicly available at https://github.com/yhhan19/variant-aware-primer-design.

## Methods

### Overview

The input to Δ-PRO is a multiple sequence alignment and optionally a reference genome. If a reference genome is not provided, then the consensus sequence of the multiple sequence alignment will be used. The first step of Δ-PRO is the estimation of heterozygosity across the genome and the identification of highly polymorphic loci. Existing methods rely on heuristic weighting schemes of SNP frequencies or entropy measures, which can overemphasize rare alleles. Here, we use Gini impurity as a more stable and biologically meaningful measure of site-based variance. Next, we use dynamic programming to select PDRs in a manner that minimizes regions that cover highly polymorphic loci. After selection of the PDRs, Primer3 is run to obtain a set of primers for each region. These primers are pooled. Finally, Δ-PRO selects a subset of primers from the pooled set that minimizes dimerization.

### Gini Impurity Computation

Gini impurity provides a simple, site-wise measure of variability in a multiple sequence alignment. For a given column, let *p*_*a*_ be the frequency of nucleotide *a*, the Gini impurity at that site *i* is

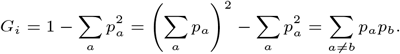

This formulation closely relates the Gini impurity to the pairwise substitution rate, providing an intuitive, functionally relevant estimate of site-wise variation. Gini impurity is reference-free, relying solely on empirical allele frequencies within a multiple sequence alignment. Moreover, it is inherently less sensitive to rare alleles, since the contribution of each allele is quadratic in its frequency, down-weighting extremely low-frequency variants unlikely to impact amplification efficiency. Finally, Gini impurity admits a simple probabilistic interpretation analogous to SNP frequency. We calculate the Gini impurity for each position in the multiple sequence alignment and store it in an array.

To illustrate the contrast between reference-based and reference-free measures of site-level variation, consider the allele frequency distributions *A* = {0.7, 0.1, 0.1, 0.1} and *B* = {0.7, 0.3}, assuming the reference allele frequency is 0.7. Both distributions yield the same SNP frequency (*f* (*A*) = *f* (*B*) = 0.30), because the SNP frequency only accounts for the total non-reference mass and therefore ignores how this mass is distributed among alternative alleles. In contrast, Shannon entropy separates the two distributions more clearly, with *H*(*A*) = 1.36 versus *H*(*B*) = 0.88, assigning a much higher value to *A* due to the presence of multiple rare alleles. The square-root–smoothed counterparts, *f* ^0.5^(*A*) = *f* ^0.5^(*B*) = 0.55, *H*^0.5^(*A*) = 1.17, and *H*^0.5^(*B*) = 0.94, are used by Olivar to reduce dynamic range and magnify small signals while still preserving the correct ordering. In practice, however, different datasets often require different smoothing exponents to balance stability and discriminability, whereas reference-free measures such as Gini impurity (*G*(*A*) = 0.48 and *G*(*B*) = 0.42) provide a more consistent and biologically interpretable assessment of site-level variation.

### PDR Optimization

Next, we use a dynamic programming algorithm to select non-overlapping PDRs from which primers are generated, aiming to avoid nucleotides with undesirable properties such as high heterogeneity. We define *rs*_*i*_ as the sequence risk associated with the *i*th nucleotide and derive *rs*_*i*_ from the previously computed Gini impurity. Importantly, our PDR optimization formulation and algorithm are agnostic to the choice of scoring function, provided that it yields a score for each nucleotide.

#### Problem Definition

Let the target genome sequence be denoted as {*g*_1_, *g*_2_, · · ·, *g*_*n*_}. For each position *i* ∈ {1, · · ·, *n*}, a score *rs*_*i*_ ∈ ℝ_≥0_ is defined according to the chosen metric (e.g., the Gini impurity can be used with *rs*_*i*_ = *G*_*i*_), where high *rs*_*i*_ values mark nucleotides that should be avoided during PDR selection. Our PDR optimization algorithm takes as input the array {*rs*_1_, *rs*_2_, …, *rs*_*n*_}. A PDR starting at position *p* is associated with a composite score *R*(*p*) computed from the segment {*rs*_*p*_, …, *rs*_*p*+*L*−1_}. The optimization task is to select a valid set of *non-overlapping* forward–reverse PDR pairs *P* = {*f*_1_, *r*_1_, …, *f*_*m*_, *r*_*m*_} that *jointly cover the target* and minimize an objective *R*(*P*). Formally, *P* is subject to the following constraints:

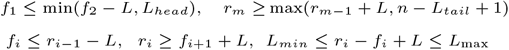

where *L*_head_ and *L*_tail_ specify the uncovered regions allowed at the beginning and end of the genome, respectively, and *L*_min_ and *L*_max_ specify the allowed range of amplicon length. We adopt the *top-α fraction* objective used in Olivar, which minimizes the aggregate score of the worst fraction of PDRs:

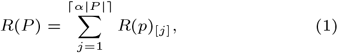

where 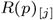 denotes the *j*th highest PDR score. This objective function generalizes several existing criteria, including the sum, min-max and fixed top-*k* formulations. This allows our algorithm to be extended to any such objective function. It also provides an explicit compromise between average-case and worst-case performance: average-case objectives may tolerate poorly performing PDRs, strict worst-case objectives can be overly conservative, and fixed top-*k* objectives may bias the solution toward selecting an unnecessarily large number of regions.

#### Exact Algorithm

We present an exact dynamic programming algorithm that explores the search space in polynomial time and ensures reproducible and optimal results. Although dynamic programming has been widely used in interval-based optimization, its application to PDR optimization is novel and non-trivial, especially under the top-*α* objective, which depends on ranking and aggregating only the highest-value elements. To avoid explicitly sorting the *R*(*p*) values, we introduce a parameter that separates the low and high *R*(*p*) values, i.e., PDRs with *R*(*p*) ≤ *u* contribute zero cost, and those with *R*(*p*) *> u* are penalized by max(0, *R*(*p*) − *u*). Varying *u* implicitly selects the worst *α* fraction of PDRs We reformulate the objective as follows.

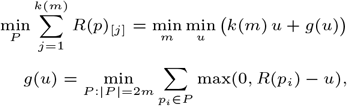

reducing the problem to a one-dimensional convex optimization over *u*. The transformed objective becomes a convex one-dimensional function of *u*, allowing the global optimum to be found efficiently by bisection, as established in Theorem 1.

##### Theorem 1

*For fixed m*, Φ_*m*_(*u*) = *k*(*m*) *u* + *g*(*u*) *is convex in u, and the global optimum u*^∗^ = arg min_*u*_ Φ_*m*_(*u*) *can be found by bisection*.

*Proof* Proof provided in the Supplementary Materials.

We now describe how to compute *g*(*u*) efficiently. The dynamic programming algorithm computes *g*(*u*) by enumerating valid PDR transitions that satisfy spacing and coverage constraints. Let opt(*i* + 1, *f*_*i*+1_, *r*_*i*+1_, *f*_*i*_, *r*_*i*_) denote the minimum risk score of a PDR set *P* consisting of *i* + 1 pairs of PDRs, where the last two pairs are (*f*_*i*+1_, *r*_*i*+1_) and (*f*_*i*_, *r*_*i*_), respectively. The recurrence is given by

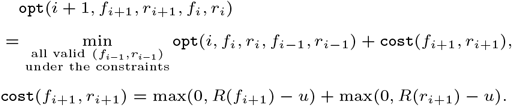

After precomputing all subproblems according to the recurrence, we derive *g*(*u*) by

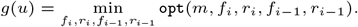

The resulting algorithm runs in 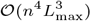 time, showing that PDR optimization is solvable in polynomial-time, assuming that a constant precision for *u* suffices. A detailed analysis of the algorithm is provided in the Supplementary Materials. With practical speedups and relaxations with approximation guarantees, our DP-based algorithm is able to produce robust PDRs in linear time. We next introduce these improvements.

#### Practical Speedups via Search Space Restriction

Although the exact algorithm is polynomial-time, its 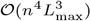 running time may be prohibitive for large genomes. We address this by introducing a restricted dynamic programming variant that substantially reduce running time while preserving the quality of the solution. We fix the distance between *f*_*i*_ and *r*_*i*_ as *r*_*i*_ − *f*_*i*_ = *L*_max_, taking advantage of the fact that the PCR amplicons are constrained to narrow length ranges. This assumption is consistent with typical primer design practice and simplifies the computation without sacrificing flexibility, since primers can still shift slightly within each PDR. Under this restriction, the number of subproblems becomes 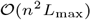, with 𝒪 (*L*_max_) transitions per state, resulting in 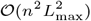 total complexity. Storing valid transitions in range-minimum-query (RMQ) structures further reduces this to 𝒪 (*n*^2^*L*_max_ log *L*_max_).

#### Relaxed Formulation with Approximation Guarantee

To achieve near-linear scaling, we relax *k*(*m*) = ⌈*α*2*m*⌉ to *k*^′^(*m*) = *α*2*m*, giving

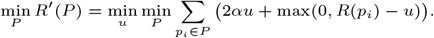

The relaxed objective remains convex and can be solved by bisection with dynamic programming in 𝒪 (*n*) time when *L*_max_ is treated as constant.

##### Theorem 2

*Let P* ^∗^ *and P* ^′^ *be the optimal PDR sets for R and R*^′^ *respectively. Then*

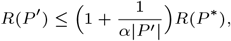

*and the approximation ratio converges to 1 as* |*P* ^′^| → ∞.

*Proof* Proof provided in the Supplementary Materials.

Theorem 2 guarantees that the relaxed near-linear solver yields PDR sets whose risk differs from the exact optimum by at most a small factor, ensuring near-optimal performance while enabling substantial speedups. In summary, we provide both an exact polynomial-time dynamic programming algorithm and a relaxed near-linear variant with theoretical guarantees. Empirical results confirm that the relaxed solution closely matches the exact optimum while scaling efficiently to genome-length targets.

### Dimerization Minimization

After computing optimal PDRs and generating primer candidates in each PDR, we devise an algorithm that selects a set of primers that minimizes dimerization. We first formulate the dimerization minimization problem as a graph optimization problem, modeling primer interactions through Δ*G*-weighted edges that represent thermodynamic compatibility, and then demonstrate how it can be solved efficiently through neighborhood search.

#### Problem Definition

We define a graph *G* = (*V, E*) whose vertices represent candidate primers: each vertex corresponds to a single primer, and the vertex set *V* is partitioned into *m* groups, one for each PDR. An edge (*u, v*) ∈ *E* is included only when *u* and *v* come from different PDRs, since exactly one primer is selected from each PDR. Each edge (*u, v*) is weighted by a dimer-prediction score reflecting the predicted interaction between the corresponding primers; specifically, we set *w*(*u, v*) = Δ*G*(*u, v*), the pre-computed Gibbs free energy for that primer pair. Our objective is to select one primer (vertex) from each PDR so that the total dimerization score (sum of edge weights) is maximized. Formally, given partitions *V*_1_, *V*_2_, · · ·, *V*_*m*_, we search for a subset of the vertices in the graph that maximizes

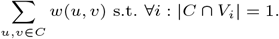

Hence, this problem is equivalent to the maximum-weight clique problem on a *m*-partite graph.

We note that more negative Δ*G* values correspond to worse dimerization, in the sense that they indicate stronger, more stable primer–primer interactions. In contrast, if a positively oriented metric for the edge weights is used—such as the badness score, where larger values correspond to worse interactions Wang et al. [2024], Xie et al. [2022]—then the objective becomes a minimum-weight clique problem. However, in either case, the underlying clique-selection problem is NP-hard and unlikely to be solved in polynomial-time Dawande et al. [2000].

#### Efficient neighborhood exploration

Due to the combinatorial complexity of the problem, we use a local search algorithm that iteratively improves the current solution by exploring its neighborhood. A neighbor is obtained by replacing exactly one primer in the current selection with an alternative candidate from the same PDR. Formally, we represent the current solution by vertices *c*_*i*_ ∈ *V*_*i*_ for each *i* ∈ 1, …, *m*, and let 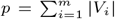 denote the total number of vertices in all partitions. A naive local search iteration considers all neighbors obtained by replacing some *c*_*j*_ with another vertex *v* ∈ *V*_*j*_, chooses the replacement that yields the largest improvement, and repeats until no further improvement is possible. Evaluating the effect of replacing *c*_*j*_ with *v* requires computing the contribution of the new vertex *v* to each of the other *m* − 1 selected vertices, which takes 𝒪 (*m*) time. Since there are 𝒪 (*p*) possible replacements, exploring the entire neighborhood would therefore require 𝒪 (*p, m*) time per iteration.

We achieve an 𝒪 (*p*)-time neighborhood scan by maintaining for every vertex *v* ∈ *V*_*j*_ the value

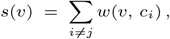

that is, the contribution of *v* to the current solution if *v* were selected instead of *c*_*j*_ . If *c*_*j*_ is replaced by some *v* ∈ *V*_*j*_ then the difference in the objective value is Δ_*j*_ (*v*) = *s*(*v*) − *s*(*c*_*j*_), which can be computed in 𝒪 (1) time. Thus, by scanning all *v* ∈ *V*_*j*_ and all *j* ∈ {1, …, *m*}, we can evaluate all 𝒪 (*p*) neighbors in 𝒪 (*p*) time and select the one yielding the best improvement. After replacing *c*_*j*_ with *v*^∗^ ∈ *V*_*j*_, we must update the values *s*(*u*) for all 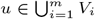. Since only one chosen vertex changes, namely *c*_*j*_ → *v*^∗^, each *s*(*u*) can be updated in 𝒪 (1) time as follows

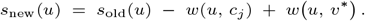

Hence, all *s*(*u*) values can be maintained in 𝒪 (*p*) total time for each iteration. This yields an 𝒪 (*p*) running time per iteration while preserving exact move evaluations.

## Experimental Results

### Experimental Setup

#### Datasets

We evaluated the performance of Δ-PRO against competing methods on two viral datasets: Foot-and-Mouth Disease (FMD) and Zika. The **FMD dataset** is a multiple sequence alignment of 146 FMD genomes, each approximately 8.4 kbp in length, but it is highly fragmented, with about 73% of positions occupied by gaps, indicating the inclusion of partial or low-coverage genomes. After filtering to retain only near-complete sequences, we obtained a refined subset of 28 genomes that span 8,180 sites with less than 1% missing data. The **ZIKA dataset** comprises the multiple sequence alignment of Zika virus genomes. The alignment contains 372 sequences that span approximately 12.2 kb, with a moderate level of missing data (approximately 12% gaps) and a well-balanced nucleotide composition. After trimming poorly aligned regions and gaps, the filtered alignment retains the same 372 sequences but is reduced to 10,753 sites, with only 13,829 gaps (*<*1.5%) and negligible ambiguity codes. Detailed information on the dataset sources and filtering criteria is provided in the Supplementary Materials.

#### Evaluation Methods

In PDR optimization, our comparison focuses on the heuristic search algorithm introduced by Olivar, as it is currently the only available method that implements a similar variant-aware design framework. In Δ-PRO, we use the relaxed optimization with length-restricted amplicon speedup for fairness and efficiency. The primer candidates of Δ-PRO are derived from PDRs using Primer3; all parameter configurations are provided in the GitHub repository. For dimerization optimization, we compare our graph-based algorithm with the SADDLE algorithm, which serves as the underlying dimerization minimization component within Olivar. In our evaluation, SADDLE was invoked through the implementation bundled within Olivar to ensure consistent parameter handling. Finally, we benchmark Δ-PRO against PrimerScheme and Olivar in an end-to-end evaluation, providing all methods with the same multiple sequence alignments and comparing the resulting primer panels in terms of design quality and robustness. In all experiments, we fixed the PDR length to 40bp, corresponding to a typical window for robust primer selection. Candidate primer pairs were generated to produce amplicons ranging from 252bp to 420bp, consistent with common tiling-PCR protocols used in viral genome sequencing. All methods were run on the HiPerGator 3.0 cluster at the University of Florida, using nodes with AMD EPYC 75F3 (Milan) processors providing 64 CPU cores at base frequency 3.0 GHz and 512 GB of RAM available per node. The core optimization algorithms were implemented in C++11 and the command line user interface was implemented in Python3.

**Table 1.** Comparison of Optimal vs. Heuristic PDR optimization across variation metrics. (A) Number of PDRs: lower values indicate more compact designs. (B) Objective function values: lower scores indicate more robust primer designs. **Improve** columns show signed relative change from Heuristic to Optimal (Improve = 1 - Optimal / Heuristic); positive values indicate improvement.

| Metrics |  | FMD |  |  | ZIKA |  |  |
| --- | --- | --- | --- | --- | --- | --- | --- |
|  |  | Optimal | Heuristic | Improve | Optimal | Heuristic | Improve |
| # of PDRs | Entropy | 25.00 ± 0.00 | 37.62 ± 1.67 | 33.5% | 36.24 ± 1.58 | 49.52 ± 0.70 | 26.8% |
|  | Entropy<sup>0.5</sup> | 24.76 ± 0.43 | 37.66 ± 1.66 | 34.2% | 34.60 ± 0.92 | 49.66 ± 0.71 | 30.3% |
|  | SNP frequency | 24.90 ± 0.30 | 37.80 ± 1.44 | 34.1% | 36.98 ± 1.38 | 49.54 ± 0.61 | 25.3% |
|  | SNP frequency<sup>0.5</sup> | 25.00 ± 0.00 | 37.68 ± 1.73 | 33.7% | 35.24 ± 1.38 | 49.42 ± 0.83 | 28.7% |
|  | Gini impurity | 24.96 ± 0.20 | 37.68 ± 1.53 | 33.8% | 36.72 ± 1.59 | 49.46 ± 0.70 | 25.8% |
| $R(P)$ Values | Entropy | 885.84 ± 67.14 | 1417.57 ± 107.95 | 37.6% | 37.18 ± 2.66 | 65.92 ± 4.20 | 43.6% |
|  | Entropy<sup>0.5</sup> | 1591.92 ± 133.50 | 2477.85 ± 150.01 | 35.8% | 136.39 ± 14.29 | 228.71 ± 14.70 | 40.4% |
|  | SNP frequency | 446.14 ± 61.58 | 637.11 ± 90.66 | 30.0% | 13.64 ± 1.91 | 25.26 ± 3.51 | 46.0% |
|  | SNP frequency<sup>0.5</sup> | 1012.00 ± 73.07 | 1571.52 ± 123.10 | 35.6% | 69.01 ± 5.90 | 119.05 ± 8.66 | 42.0% |
|  | Gini impurity | 674.85 ± 61.44 | 1017.87 ± 102.95 | 33.7% | 28.13 ± 2.47 | 51.42 ± 4.44 | 45.3% |

### PDR Optimization

Our algorithm consistently yields fewer PDRs and lower objective function values than the heuristic method, as shown in Figure 1. Across all variation metrics, the optimal PDR algorithm reduces the number of PDRs by approximately 33% on the FMD dataset and 28% on the ZIKA dataset, yielding more compact design regions. The corresponding *R*(*P*) values decrease by roughly 35% on FMD and 38% on ZIKA, indicating substantially improved robustness. This reduction reflects the ability of the exact optimization procedure to eliminate redundant or overlapping regions that arise in heuristic sampling. The optimal solver directly maximizes coverage and robustness under explicit combinatorial constraints, ensuring that each selected PDR contributes meaningfully to the overall coverage of the target. In contrast, heuristic sampling may select partially redundant regions that inflate the total count without improving variant robustness. Using fewer PDRs also offers practical advantages, including reduced primer count, lower risk of cross-dimerization, simpler pooling, and overall improved robustness of multiplex PCR workflows.

**Fig. 1.**
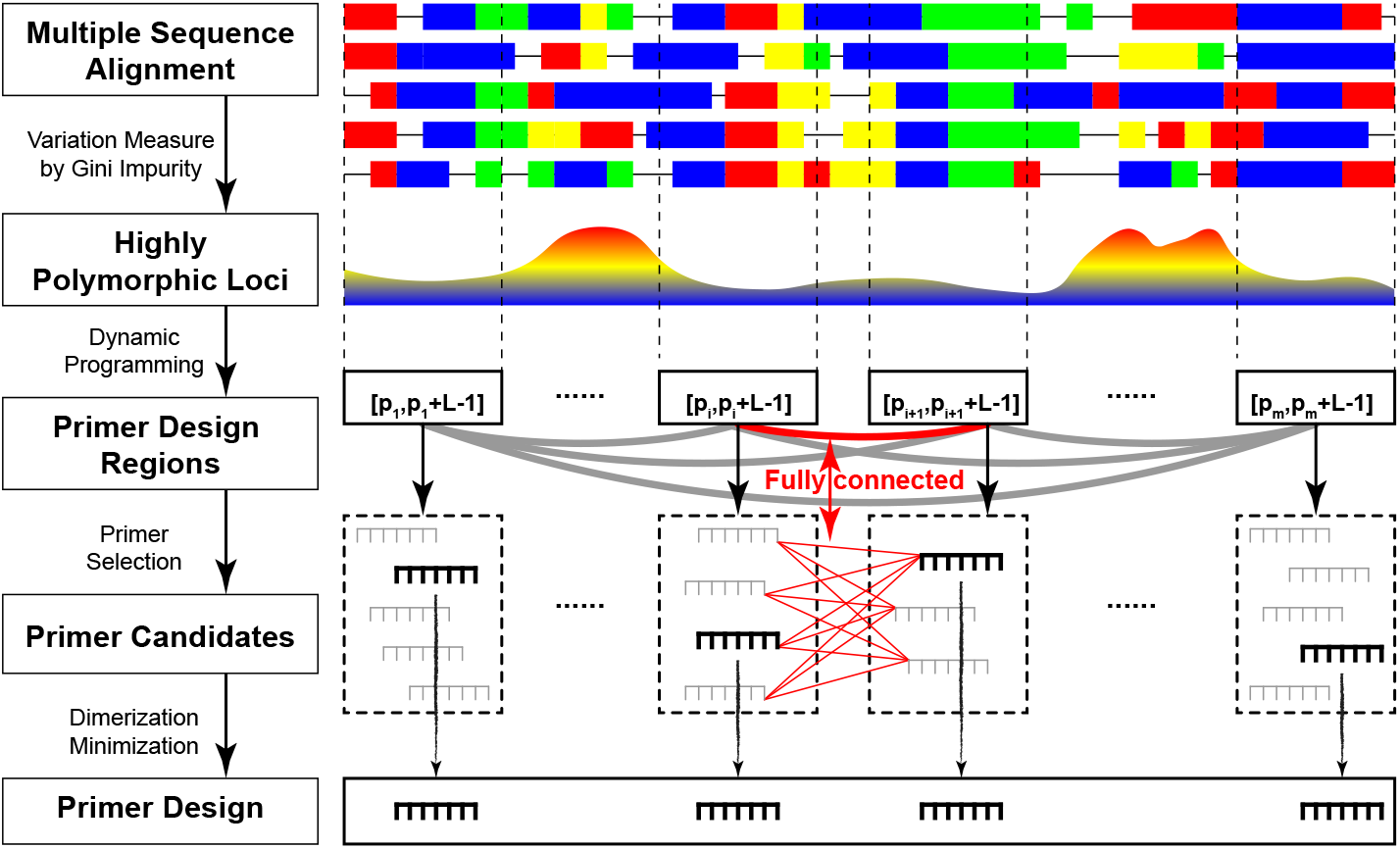
Starting from a multiple sequence alignment, Δ-PRO computes a nucleotide-level variation profile using Gini impurity and applies dynamic programming to select low-risk PDRs under amplicon-length constraints. Candidate primers generated from each PDR are then represented as vertices in a *k*-partite graph with edges weighted by predicted Δ*G*, and a local-search heuristic selects one primer per PDR to minimize primer–primer dimerization while maintaining target coverage. For visual clarity, we omit the fully connected edges among vertices from non-adjacent PDRs.

Although the *R*(*P*) objective can flexibly incorporate diverse site-level metrics, it is not obvious that minimizing this objective alone will always produce biologically robust regions. Therefore, rather than comparing objective values, we evaluate each method by examining the resulting PDRs and measuring their robustness to sequence variation. For an MSA with *N* sequences and a PDR spanning coordinates [*p, p* + *L* − 1], we define its mismatch score as the average number of positions where sequences differ from the reference:

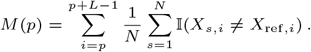

This reflects the expected mismatch rate for primers derived from that region. Mismatch scores are computed for all PDRs, ignoring gaps and ambiguous bases. To compare methods, we focus on the most variable regions—the top-10 (absolute) or top-10% (relative) highest-mismatch PDRs—as these represent the most failure-prone sites for downstream PCR. Lower mismatch counts in this upper tail indicate more variant-robust primer designs.

Figure 2 summarizes the results for both the FMD and Zika datasets under the two optimization strategies (optimal and heuristic). The optimal strategy consistently yields fewer mismatches, confirming the benefit of exact optimization. Relative measures (top-10%) show tighter distributions due to normalization, while absolute comparisons demonstrate clear improvements even when the number of regions is fixed. Notably, even though the optimal method often selects fewer PDRs—a more constrained setting with less flexibility to maintain coverage—its highest-mismatch regions still exhibit fewer mismatches than those produced by the heuristic approach. The FMD dataset exhibits greater variability due to greater genomic diversity, whereas ZIKA shows narrower distributions consistent with lower divergence. Square-root transformations improve robustness for FMD but degrade it for ZIKA, suggesting over-smoothing in low-diversity alignments– —a scenario that is more common in practice. By contrast, the Gini metric performs reliably across both datasets, producing stable and variant-robust PDRs.

**Fig. 2.**
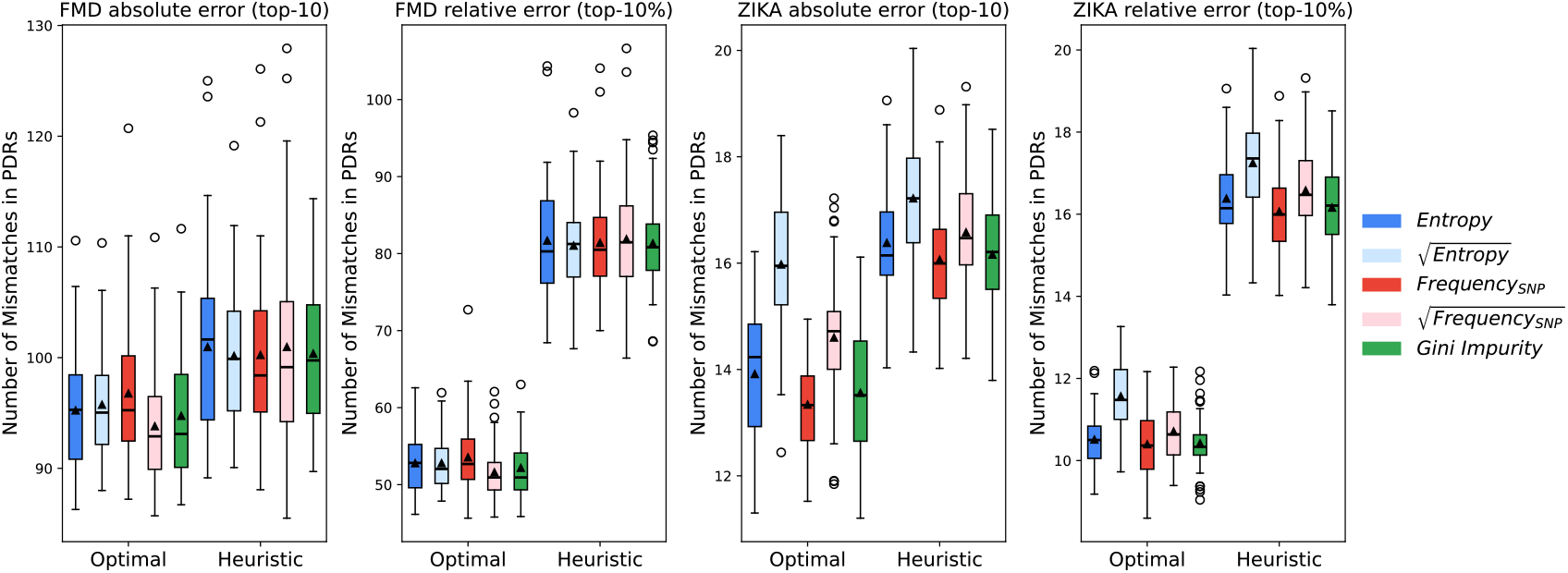
Comparison of mismatches in PDRs for FMD and ZIKA datasets. Each boxplot summarizes the total number of mismatches observed in candidate PDRs generated under different optimization methods as well as variation metrics. The y-axis denotes the number of mismatches in PDRs, while the x-axis separates results obtained from the optimal and heuristic PDR optimization methods. The left and right subplots of each dataset correspond respectively to: (1) absolute error— total mismatches in the ten most variable PDRs for the FMD dataset; (2) relative error—mismatches normalized by the total number of PDRs (top 10% subset). Colored boxes represent different variant metrics, including empirical entropy and SNP frequency, as well as their square roots (suggested by Olivar), and Gini impurity.

Across both datasets, our dynamic programming algorithm for optimal PDR optimization is substantially faster than the Olivar heuristic. For FMD, the dynamic programming running time is approximately 6 seconds on average, compared to 94 seconds for the heuristic. On the ZIKA dataset, the gap is even larger: 8.6 seconds versus 154 seconds. This empirical speedup is consistent with the asymptotic behavior: the DP algorithm runs in 𝒪 (*n*) time in the genome length *n*, whereas the Olivar heuristic is 𝒪 (*n*^2^) because it performs 𝒪(*n*) iterations, each requiring 𝒪(*n*) time for random PDR generation. Both methods exhibit linear memory growth with genome length *n*, but the DP approach has a larger constant factor. We also report the memory usage of the DP algorithm, as it must store intermediate states and therefore consumes substantially more memory, making it the primary memory bottleneck of Δ-PRO. On FMD, DP requires 328 MB compared to 131.5 MB for the heuristic, and on the larger ZIKA dataset it uses 501.5 MB versus 243.5 MB.

### Dimerization Minimization

We compare our approach with SADDLE to assess its effectiveness in minimizing primer dimerization. A common set of primer candidates is generated first from the PDRs identified by Olivar, using identical input parameters across all methods. Each method is then applied to this shared candidate pool to produce a final primer set, ensuring a fair comparison. This procedure is repeated 10 times with different random seeds, yielding 10 distinct PDR sets and 20 primer pools for each dataset.

For Δ*G* values, Δ-PRO consistently achieves more favorable scores in all replicates, as SADDLE does not directly optimize this objective. In addition, we evaluate performance using the *Badness score* so that both methods take as input identical primer candidates and dimer predictions. Across 20 pools of FMD data, Δ-PRO achieves lower Badness scores in 19 cases and matches SADDLE in the remaining pool. On average, SADDLE obtains a Badness score of 682.50, while Δ-PRO reduces this to 662.87. On the Zika dataset, SADDLE produces an average badness score of 915.45, whereas Δ-PRO achieves a lower mean of 864.08. Across 20 pools, our method outperforms in 16 cases and ties in the remaining 4, demonstrating consistently improved performance. The results show that the graph-based algorithm outperforms the simulated-annealing approach employed by SADDLE. In Figure 3, we present the predicted dimerization distributions for the primer pairs generated by the two methods, using the input data based on the FMD dataset and produced by the first random seed as an example. Across both pools and across both evaluation objectives, Δ-PRO yields distributions that are shifted toward better designs with reduced predicted dimerization.

**Fig. 3.**
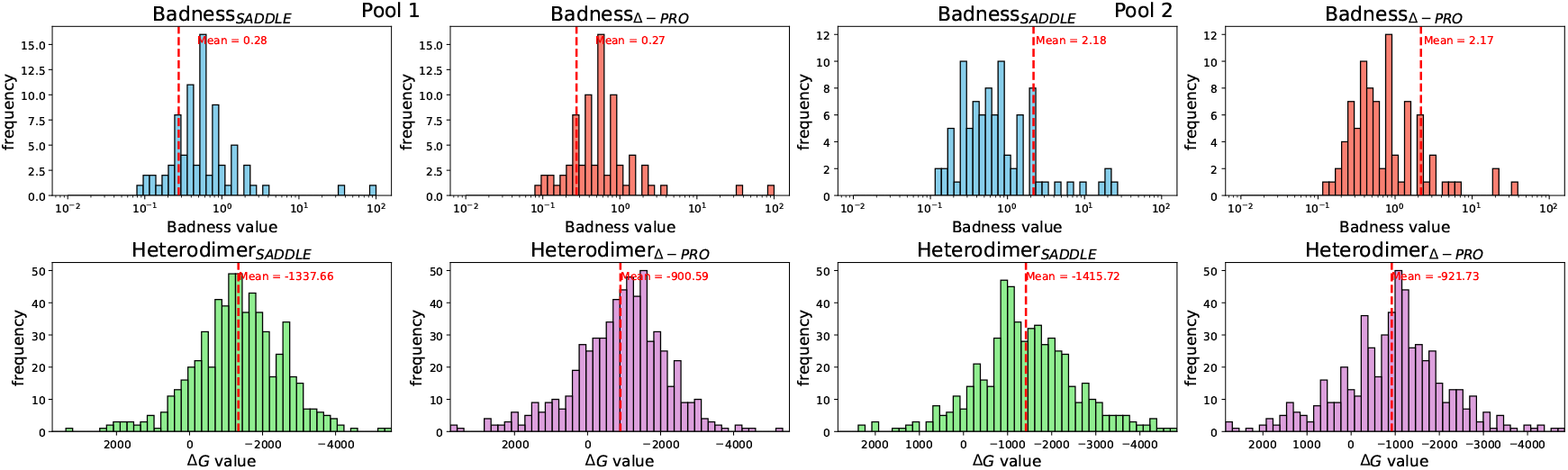
Example histograms of badness scores (top) and heterodimer Δ*G* values (bottom) for primer sets selected by SADDLE and our Δ–PRO optimization, shown for Pool 1 and Pool 2. These results correspond to PDRs generated using a fixed random seed (127978094) for reproducibility. Note that most badness values are 0 and therefore do not appear in the log-scale plots. The complete results for all replicates are available in the Supplementary Materials.

On the FMD dataset, SADDLE requires about 1.5 minutes per run (10 replicates), while Δ-PRO (iteration limit 10^4^) finishes in roughly 34 seconds on average, with the core graph-optimization taking only 3.5 seconds and most of the runtime spent constructing the graph and invoking the Olivar-based Badness function. On the ZIKA dataset, SADDLE requires about 2.5 minutes per run, while Δ-PRO runs fast with an average of 22 seconds of which roughly 4.1 seconds are spent in the optimization step. Overall, our method outperforms SADDLE not only in solution quality but also in computational efficiency across both datasets.

### End-to-End Benchmarking

Finally, we present an end-to-end evaluation that compares our approach with PrimalScheme and Olivar. We construct simulation datasets based on the real FMD and ZIKA multiple sequence alignment to accommodate all methods, since existing tools often impose distinct assumptions or requirements on the input data. We use the consensus genomes derived from the FMD and ZIKA multiple sequence alignments as references, and we generate synthetic variants by following the simulation procedure described in Wang et al. [Wang et al. [2024]. For the FMD dataset, we generate 28 simulated sequences to reflect the size of the real MSA. For the ZIKA dataset, the simulation is limited to 100 sequences due to restrictions in PrimalScheme. Subsequently, we apply each method to the simulated datasets and evaluate the resulting primers using multiple metrics, including sequence mismatches and predicted dimerization.

Across both FMD and ZIKA datasets, Δ-PRO achieves competitive or superior robustness to sequence variation while using substantially fewer primer pairs than Olivar and PrimalScheme. For fair comparison, in addition to reporting Top-10% and total mismatches across primer sets of different sizes, we use the Top-*k*_*m*_ metric, which evaluates only the *k*_*m*_ worst primer pairs in each panel. Here, *k*_*m*_ is set to the number of primer pairs produced by Δ-PRO, which generates the smallest panel among the three methods. Even under equal panel sizes, Δ-PRO achieves lower mismatch counts — whereas the extra primer pairs generated by the other methods only introduce additional mismatch burden. Moreover, Δ-PRO consistently yields predicted dimerization energies closer to zero than competing methods, suggesting a reduced risk of strong primer–dimer formation. Overall, Δ-PRO delivers a favorable balance of robustness, compact primer panels, and improved dimerization profiles compared with existing methods. Δ-PRO is also more efficient, with end-to-end runtimes of 98.4 seconds on FMD and 188.6 seconds on ZIKA, while Olivar and PrimalScheme require considerably more time on both datasets. Using an 8-thread parallel setup, Olivar required 6.3 minutes (FMD) and 7.7 minutes (ZIKA), longer than Δ-PRO on both datasets. PrimalScheme was substantially slower, taking 9 minutes on FMD and 29 minutes on ZIKA.

**Table 2.** End-to-end evaluation of primer sets from different methods on FMD and ZIKA datasets. Lower mismatch values indicate more robust primers, and more negative Δ*G* values indicate stronger predicted dimerization. The metric Top-*k*_*m*_ denotes the truncated mismatch sum, computed by restricting each method to the same number of primer pairs as used by Δ-PRO, ensuring a fair comparison across methods with different primer counts.

| Dataset | Method | #Pairs | Variant Mismatches |  |  | Pool 1 Dimerization |  | Pool 2 Dimerization |  |
| --- | --- | --- | --- | --- | --- | --- | --- | --- | --- |
| | | | Top-$k_m$ | Top-10% | Total | Mean $\Delta G \pm \text{Std}$ | | Mean $\Delta G \pm \text{Std}$ | |
| FMD | Δ-PRO | 25 | 138.21 | 30.00 | 138.21 | **-798.23 ± 1373.62** | | **-970.88 ± 1187.39** | |
| | Olivar | 38 | 155.68 | 37.43 | 186.61 | -1404.59 ± 1250.60 | | -1404.58 ± 1089.36 | |
| | PrimalScheme | 38 | 157.21 | 41.71 | 180.43 | -1276.06 ± 1136.64 | | -1269.63 ± 1174.80 | |
| ZIKA | Δ-PRO | 38 | 39.30 | 8.33 | 39.30 | **-1096.60 ± 1192.61** | | **-1073.27 ± 1228.11** | |
| | Olivar | 49 | 47.80 | 11.67 | 49.87 | -1267.81 ± 1143.81 | | -1319.77 ± 1082.96 | |
| | PrimalScheme | 54 | 42.81 | 10.68 | 47.22 | -1302.50 ± 1141.37 | | -1262.53 ± 1102.85 | |

## Conclusion

In this work, we introduced Δ-PRO, a scalable framework for variant-aware multiplex primer design that jointly optimizes where to place primers and which primers to select. By combining a biologically interpretable variation metric based on Gini impurity with a dynamic programming formulation of the PDR optimization problem, Δ-PRO yields PDR sets that are both more compact and more robust to sequence variation than those produced by existing heuristic approaches. Across FMD and ZIKA viral genomes, our exact and relaxed solvers consistently reduced the number of PDRs while lowering mismatch burdens in the most variable regions, indicating that principled optimization produces designs better aligned with downstream amplification robustness. Moreover, Δ-PRO enables explicit modeling of primer–primer interactions at the scale required for modern multiplex panels, where naive enumeration is computationally infeasible. Future work includes extending Δ-PRO to jointly optimize on-target coverage, off-target binding, and dimerization within a single objective function, which would further broaden its applicability, while integration with experimental validation will be essential to demonstrate that it can make large-scale, variant-aware multiplex PCR design more reliable and routine in both research and surveillance settings.

