## Supplemental Materials for "Scaling Variant-Aware Multiplex Primer Design"

### Supplementary Materials for
### Scaling Variant-Aware Multiplex Primer Design

#### 1 PDR Optimization Algorithm Details

##### 1.1 Problem Formulation

In this section, we introduce a novel dynamic programming (DP)-based approach for optimizing Primer Design Regions (PDRs). PDRs are genomic regions from which primer candidates are generated as contiguous subsequences. We begin with the formulation of the optimization problem from `Olivar`. `Olivar` quantifies undesired sequence features by assigning per-nucleotide risk scores across the genome. The resulting risk array is simply a vector aligned to the target genome, where each entry corresponds to the "risk" of using that nucleotide in primer design (so a PDR overlaps it). Each base position is evaluated for properties that could lead to poor performance in multiplex PCR, for example, regions with heavy variations and secondary structures, low sequence complexity, extreme GC content, and non-specificity. In practice, the risk array is not tied to a strict definition; instead, it can be computed from a variety of information sources depending on the availability of the data; for example, `Olivar` computes the risk score of a position as linear combination of contributions from SNP frequency, extreme GC content, sequence complexity, and non-specificity: $rs_i = w_1 \cdot \text{snp}(i) + w_2 \cdot \text{egc}(i) + w_3 \cdot \text{lc}(i) + w_4 \cdot \text{ns}(i)$, where $w_1$, $w_2$, $w_3$, and $w_4$ are user-defined weights (set to 1 by default). We discuss our approach for improving the computation of risk scores in the main text, and focus here on the PDR optimization problem.

**Problem Definition.** Let the target genome sequence be denoted as $G = \{g_1, g_2, \cdots, g_n\}$. For each position $i \in \{1, \cdots, n\}$, a risk score $rs_i \in \mathbb{R}_{\geq 0}$ is defined. Larger values of $rs_i$ indicate higher risk for any PDR overlapping position $i$, thereby reducing the suitability of the corresponding region for primer design. A PDR of length $L$ is a contiguous subsequence of $G$ spanning positions $p$ through $p+L-1$. In the following, we assume that all PDRs have the same fixed length $L$, and represent each PDR by its starting position. The risk score of a PDR starting at position $p$ is defined as a function of the per-nucleotide risks within its interval:

$$R(p) : \{p \mid 0 \leq p \leq n - L + 1\} \mapsto \mathbb{R}_{\geq 0},$$

where $R(p)$ may be computed based on the risk scores in the PDR: $rs_p$, $\cdots$, $rs_{p+L-1}$. Given a set of PDR design requirements — e.g., target coverage, multiplexing constraints, or a maximum number of regions — the PDR optimization problem is to select a collection of non-overlapping PDRs such that a total risk objective is minimized. In order to ensure that the amplicons (defined by pairs of forward and reverse primers) collectively cover all desired loci of the target, PDRs are also generated in pairs. Let the set of PDRs be $P = \{f_1, r_1, f_2, r_2, \cdots, f_m, r_m\}$, where $m$ is the number of pairs in $P$, and $f_i$ and $r_i$ are the starting positions of the forward and reverse PDRs corresponding to the $i$-th amplicon. We require the PDRs in $P$ to satisfy the following constraints for all $i$:

$$f_1 \leq \min(f_2 - L, L_{head}), \quad r_m \geq \max(r_{m-1} + L, n - L_{tail} + 1)$$
$$f_i \leq r_{i-1} - L, \;\; r_i \geq f_{i+1} + L, \;\; L_{min} \leq r_i - f_i + L \leq L_{\max}$$

These inequalities ensure that forward and reverse PDRs are placed so that amplicons are properly defined, do not overlap improperly, and collectively cover the entire target sequence, leaving at most $L_{head}$ bases uncovered at the beginning and $L_{tail}$ bases uncovered at the end. In addition, the user may impose bounds on amplicon length, requiring it to lie within a prescribed interval $[L_{min}, L_{\max}]$.

**Optimization Objectives.** Different formulations of the PDR optimization objective can emphasize distinct aspects of design robustness. A straightforward choice is the *sum objective*, which minimizes total risk but may neglect high-risk regions. In contrast, the *min–max objective* instead focuses on controlling the worst-case risk, ensuring that no single PDR exhibits an excessively high score, though this can lead to overly conservative designs. The *top-k objective* aggregates only the $k$ highest-risk PDRs to prioritize minimizing the most problematic regions without penalizing low-risk ones. A natural generalization is the *top-$\alpha$ fraction objective*, adopted in `Olivar`, which considers a fixed proportion $\alpha$ of the riskiest PDRs, scaling adaptively with the total number of amplicons.

Our method is general and supports all the objective formulations introduced above. Among these, the top-$\alpha$ fraction objective is the most challenging, as it requires ranking and aggregating only the riskiest subset of PDRs, introducing nonlinearity into the optimization landscape. Nevertheless, it provides a flexible trade-off between minimizing average and worst-case risks, and thus serves as our focus in the following sections. Accordingly, we adopt the top-$\alpha$ fraction formulation in the following algorithms. Under this objective, the total risk of a valid PDR set is evaluated by aggregating the top-$k$ highest-risk PDRs, where $k = \lceil \alpha \cdot |P| \rceil$ corresponds to the top-$\alpha$ fraction of all PDRs:

$$R(P) = \sum_{j=1}^{k} R(p)_{[j]},$$

where $R(p)_{[j]}$ denotes the $j$-th greatest risk score among all PDRs in $P$. Therefore, only the $k$ worst-scoring regions contribute to the loss, allowing the objective to emphasize the highest-risk components of the design. In `Olivar`, the parameter $k$ is set to $k = \lceil m/5 \rceil$ by default so only top 10% of PDRs with the highest risk scores contribute to the loss. In practice, the user may specify the proportion of highest-risk PDRs to consider by a function $k(m) = \lceil \alpha \cdot 2m \rceil$ of the total number of PDRs in $P$, where $\alpha \in (0, 1]$.

##### 1.1.1 Exact Optimization

`Olivar` addresses PDR optimization using a heuristic that repeatedly samples random PDR sets and keeps the one with the lowest risk. However, this approach lacks optimality guarantees, scales poorly with problem size, and produces variable results across runs. We instead propose a DP–based method that explores the solution space efficiently in polynomial time while ensuring reproducible and near-optimal solutions. Although dynamic programming has been widely applied to interval-based optimization problems, it has not previously been used for PDR optimization. Moreover, solving the top-$\alpha$ fraction objective with DP is non-trivial because it requires ranking and selectively aggregating only the highest-risk regions. Our method reformulates the discrete risk minimization into a convex form, enabling efficient one-dimensional optimization and a structured DP solution.

We reformulate the PDR selection problem into a DP structure that allows efficient computation of the optimal PDR set $P^* = \arg\min_P R(P)$. We show that $P^*$ can be computed efficiently by reformulating the objective as follows:

$$\begin{aligned}
\min_P \sum_{j=1}^{k(m)} R(p)_{[j]} &= \min_P \min_u \big(k(m)\cdot u + \sum_{p_i\in P} \max(0, R(p_i)-u)\big) \\
&= \min_m \min_{P:|P|=2m} \min_u \big(k(m)\cdot u + \sum_{p_i\in P} \max(0, R(p_i)-u)\big) \\
&= \min_m \min_u \big(k(m)\cdot u + \min_{P:|P|=2m} \sum_{p_i\in P} \max(0, R(p_i)-u)\big) \\
&= \min_m \min_u \big(k(m)\cdot u + g(u)\big).
\end{aligned}$$

The optimization objective becomes $\min_m \min_u \Phi_m(u)$ after the reformulation, where $\Phi_m(u) = k(m)\cdot u + g(u)$. We minimize the one-dimensional objective $\min_u \Phi_m(u)$ for each $m$. Having reduced the problem to a one-dimensional minimization over $u$, we next establish the convexity of the reformulated objective, which guarantees that its global optimum $u^* = \arg\min_u \Phi_m(u)$ can be efficiently found by bisection. The convexity of the reformulated objective follows from the structure of $\Phi_m(u)$, as shown in the theorem below.

**Theorem 1.1.** *For a fixed $m$, $\Phi_m(u)$ is convex in $u$, so a global optimum $u^* \in \arg\min_u \Phi_m(u)$ exists and can be found by bisection.*

*Proof.* For any feasible PDR set $P$, define

$$g_P(u) = \sum_{p_i\in P} \max(0, R(p_i)-u).$$

Each term $\max(0, R(p_i)-u)$ is convex and piecewise linear in $u$, hence $g_P(u)$ is convex and piecewise linear. The function $g(u)$ is defined as the pointwise minimum of $\{g_P(u)\}$ across all feasible sets $P$. The pointwise minimum of convex functions is convex, so $g(u)$ is convex and piecewise linear. Adding the linear term $k(m)\cdot u$ preserves convexity, establishing that $\Phi_m(u)$ is convex and piecewise linear. Because $\Phi_m(u) \to \infty$ as $u \to \pm\infty$, a global minimizer $u^*$ exists. Furthermore, since $\Phi_m(u)$ is one-dimensional and convex, its minimum can be located efficiently by bisection on the sign of its (sub)gradient, guaranteeing convergence to the global optimum. □

Finally, we describe how $g(u)$ can be computed efficiently using dynamic programming. Let $\texttt{opt}(i+1, f_{i+1}, r_{i+1}, f_i, r_i)$ denote the minimum risk score of a PDR set $P$ consisting of $i+1$ pairs of PDRs, where the last two pairs are $(f_{i+1}, r_{i+1})$ and $(f_i, r_i)$, respectively. The value of $\texttt{opt}(i+1, f_{i+1}, r_{i+1}, f_i, r_i)$ can be obtained by enumerating all feasible preceding pairs $(f_{i-1}, r_{i-1})$, selecting the optimal transition, and then adding the contribution of $(f_{i+1}, r_{i+1})$:

$$\texttt{opt}(i+1, f_{i+1}, r_{i+1}, f_i, r_i) = \min_{\substack{\text{all valid } (f_{i-1}, r_{i-1}) \\ \text{under the constraints}}} \texttt{opt}(i, f_i, r_i, f_{i-1}, r_{i-1}) + \texttt{cost}(f_{i+1}, r_{i+1})$$

$$\texttt{cost}(f_{i+1}, r_{i+1}) = \max(0, R(f_{i+1}) - u) + \max(0, R(r_{i+1}) - u)$$

After precomputing all DP subproblems according to the recurrence, we derive $g(u)$ by

$$g(u) = \min_{f_i, r_i, f_{i-1}, r_{i-1}} \texttt{opt}(m, f_i, r_i, f_{i-1}, r_{i-1}),$$

in which we enumerate all possible PDR set size $2m$.

**Complexity Analysis.** The number of subproblems is $O(n^5)$, and each subproblem requires $O(n^2)$ transitions. Thus, the total cost of filling the DP table is $O(n^7)$, which is polynomial. In practice, the number of preceding pairs of PDRs is limited by $L_{\max}$, so the number of subproblems reduces to $O(n^3 L_{\max}^2)$ while the number of transitions reduces to $O(nL_{\max})$, and therefore the complexity is $O(n^4 L_{\max}^3)$. After this preprocessing, we enumerate at most $O(n)$ possible values of $m$. Assuming the bisection search for $u$ terminates at a constant precision, the computation of $g(u)$ requires $O(n^2 L_{\max}^2)$ time by enumerating all values of $\texttt{opt}(m, f_i, r_i, f_{i-1}, r_{i-1})$. Therefore, obtaining the optimal solution $P^*$ has a complexity of $O(n^3 L_{\max}^2)$. Combining both phases, the overall time complexity of the algorithm is $O(n^4 L_{\max}^3)$, and thus polynomial under the assumption that a constant precision for $u$ is sufficient to produce an optimal solution $P^*$.

This demonstrates that, under realistic constraints and even the complex top-$\alpha$ fraction objective, the PDR optimization problem is polynomial-time solvable rather than NP-hard. Any heuristic (like the random-sampling strategy in `Olivar`) is not theoretically necessary. By showing that PDR optimization admits an exact polynomial-time solution, we establish its tractability and provide the first principled alternative to heuristic sampling approaches used in prior tools.

**Practical Speedups.** Although we have shown that the optimization problem can be solved in polynomial time, the $O(n^4 L_{\max}^3)$ runtime remains impractical for large genomes. We therefore introduce two practical variants that reduce the search space and number of subproblems while preserving solution quality. In addition, we employ data structures that allow efficient retrieval of optimal transitions. Empirically, the two variants achieve similar performance and both outperform the `Olivar` heuristic while achieving substantial speedups. In practice, the appropriate variant can be selected according to experimental requirements.

The first speedup fixes the distance between $f_i$ and $r_i$ as $r_i - f_i = L_{\max}$, exploiting the fact that amplicons in standard PCR protocols are constrained to narrow length ranges. This assumption is biologically reasonable, as PCR amplification is most reliable within a narrow length range. Fixing the inter-PDR distance therefore reflects standard laboratory practice while simplifying computation. Note that actual amplicon lengths may still vary slightly, since primers can shift within each PDR, preserving biological flexibility without compromising efficiency. Under this restriction, the number of subproblems becomes $O(n^2 L_{\max})$, and the transition cost reduces to $O(L_{\max})$, giving a total complexity of $O(n^2 L_{\max}^2)$. We further store valid transitions for each $r_i$ in data structures that support range minimum queries (RMQ) on $r_{i-1}$, reducing the transition cost to $O(\log L_{\max})$ with $O(L_{\max})$ preprocessing. This yields an overall complexity of $O(n^2 L_{\max} \log L_{\max})$.

Alternatively, we may constrain the search space by requiring that the start of the next PDR pair lies within the second half of the current one, thereby eliminating the need to consider order-two dependence in the DP recurrence. This idea builds on the observation that amplicons in multiplex PCR generally do not overlap extensively—an assumption also adopted by the primer design method `varVAMP` to simplify compatibility constraints. Under this restriction, the number of subproblems becomes $O(n^2 L_{\max})$ and the number of transitions $O(L_{\max}^2)$, giving an overall complexity of $O(n^2 L_{\max}^3)$. The RMQ-based data structure remains applicable—albeit with higher complexity due to the quadratic transitions—further reducing the runtime to $O(n^2 L_{\max} \log^2 L_{\max})$.

While these optimizations substantially improve runtime, the algorithm still scales quadratically with sequence length. To achieve near-linear scaling, we next introduce a relaxed formulation that removes dependence on the number of PDRs, attaining linear time complexity when $L_{\max}$ is treated as constant, while maintaining provable approximation guarantees. As typical amplicon lengths are only a few hundred bases, this assumption holds in practice, allowing the algorithm to scale efficiently to long genomic targets.

##### 1.1.2 Relaxed Formulation with Approximation Guarantee

To this end, we propose the following relaxed formulation, which achieves linear-time performance with respect to the sequence length while preserving a bounded approximation ratio that converges to one as $n$ increases. We relax the optimization objective by replacing $k(m) = \lceil \alpha \cdot 2m \rceil$ with $k'(m) = \alpha \cdot 2m$:

$$\min_P R'(P) = \min_u \min_P \sum_{p_i \in P} 2\alpha \cdot u + \max(0, R(p_i) - u) = \min_u g'(u),$$

where $g'(u)$ is still convex and can therefore be solved efficiently using DP (by adding $2\alpha \cdot u$ to the $\texttt{cost}(\cdot)$ function). Consequently, we can compute the relaxed optimum by performing a bisection search over $g'(u)$. The following theorem establishes a rigorous theoretical guarantee for the relaxed formulation, demonstrating that the proposed heuristic achieves near-optimal accuracy even on large genomes.

**Theorem 1.2.** *Let $P^* \in \arg\min_P R(P)$ be an optimal PDR for the original objective and $P' \in \arg\min_P R'(P)$ be an optimal PDR for the relaxed objective. Then*

$$R(P') \ \leq \ \left(1 + \frac{1}{\alpha \cdot |P'|}\right) \cdot R(P^*).$$

*Proof.* For $u \geq 0$, define

$$R(P; u) = \lceil \alpha |P| \rceil \cdot u + \sum_{p_i \in P} \max(0, R(p_i) - u), \ \ R'(P; u) = \alpha |P| \cdot u + \sum_{p_i \in P} \max(0, R(p_i) - u).$$

And we have $R(P) = \min_u R(P;u)$ and $R'(P) = \min_u R'(P;u)$. Note $R'(P;u) \leq R(P;u)$, i.e., the relaxed function has a smaller value for all $(P,u)$. Assume $(P^*, u^*)$ minimizes $R(P;u)$ while $(P', u')$ minimizes $R'(P;u)$. By definition of $R'$, we have

$$R'(P';u') = \alpha|P'| \cdot u' + \sum_{p_i \in P'} \max(0, R(p_i) - u') \quad \geq \quad \alpha|P'| \cdot u'.$$

Since $(P', u')$ minimizes $R'$, it follows that

$$R'(P';u') \leq R'(P^*;u^*) \leq R(P^*;u^*).$$

Thus

$$u' \leq \frac{R'(P';u')}{\alpha|P'|} \leq \frac{R(P^*;u^*)}{\alpha|P'|}.$$

Moreover, for any $(P,u)$, we have

$$R(P;u) - R'(P;u) = (\lceil \alpha|P| \rceil - \alpha|P|) \cdot u < u,$$

so in particular

$$R(P';u') \leq R'(P';u') + u' \leq R(P^*;u^*) + u'.$$

Combining with the bound on $u'$ gives

$$R(P';u') \leq R(P^*;u^*) + \frac{R(P^*;u^*)}{\alpha|P'|} = \Big(1 + \frac{1}{\alpha|P'|}\Big) \cdot R(P^*;u^*).$$

Finally, since $R(P) = \min_u R(P;u)$, we have $R(P') \leq R(P';u')$ and $R(P^*) = R(P^*;u^*)$, which gives the claim. If the amplicon length is bounded by a constant $L_{\max}$, then $|P'| = \Theta(n)$ as the sequence length $n$ grows. Hence the factor $1 + 1/(\alpha|P'|) \to 1$ as $n \to \infty$. □

In summary, our method computes the exact optimum in polynomial time, and its relaxed variant achieves near-linear complexity with asymptotically optimal accuracy. Experiments confirm that the relaxed solution remains close to the exact optimum in practice.

### 2 Supplementary Results

#### 2.1 Data Preparation

The FMD dataset consists of multiple sequence alignments of Foot-and-Mouth Disease Virus (FMDV) genomes. The raw alignment contains 146 sequences of approximately 8.4 kb, but is highly fragmented, with about 73% of positions occupied by gaps, indicating the inclusion of partial or low-coverage genomes. After filtering to retain only near-complete sequences, a refined subset of 28 genomes remains, spanning 8,180 sites with less than 1% missing data. This cleaned dataset exhibits balanced nucleotide composition (A, C, G, and T in comparable proportions) and minimal ambiguity, making it well-suited for computing site-based variation metrics. The ZIKA dataset comprises multiple sequence alignments of Zika virus genomes. The raw alignment contains 372 sequences spanning approximately 12.2 kb,

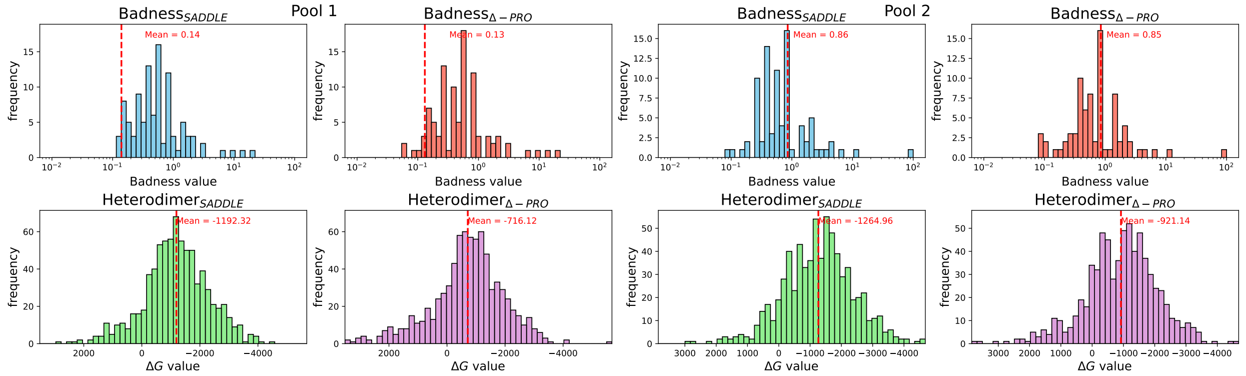


Figure 1: Results for primer candidates generated by Olivar (random seed `1181241943`) on the FMD dataset.

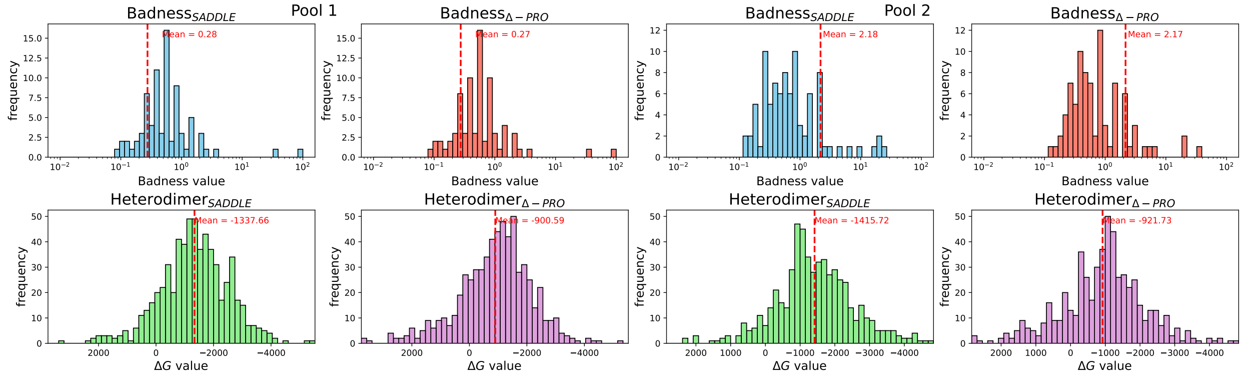


Figure 2: Results for primer candidates generated by Olivar (random seed `127978094`) on the FMD dataset.

with a moderate level of missing data—about 546,777 gap characters (12%)—and a well-balanced nucleotide composition. After trimming poorly aligned or gappy regions, the filtered alignment retains the same 372 sequences but is reduced to 10,753 sites, with only 13,829 gaps (1.5%) and negligible ambiguity codes (R, Y, N, etc.). This refined dataset captures the full genetic diversity of circulating Zika virus lineages while ensuring high alignment quality and near-complete genome coverage. The raw and filtered MSA data is available at `https://github.com/yhhan19/variant-aware-primer-design`.

#### 2.2 Histogram Distributions of Dimerization Metrics on Simulated Data

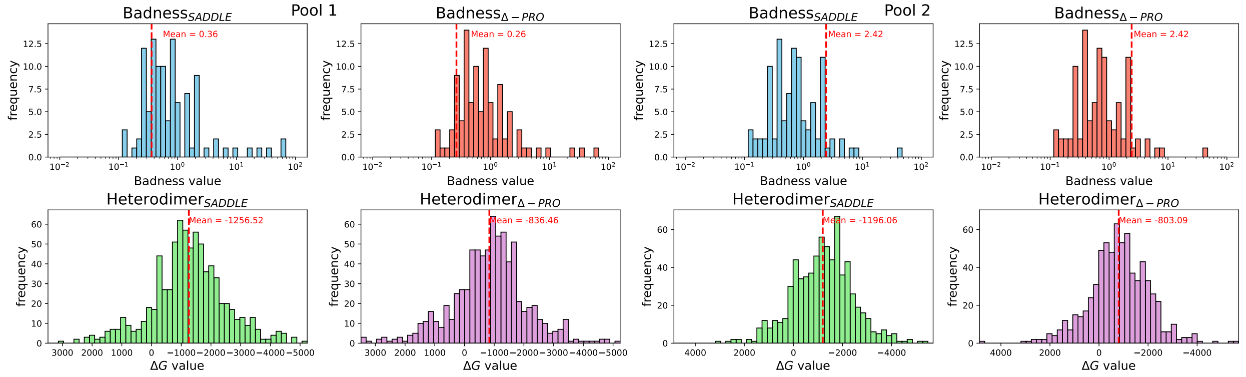


Figure 3: Results for primer candidates generated by Olivar (random seed 1812140441) on the FMD dataset.

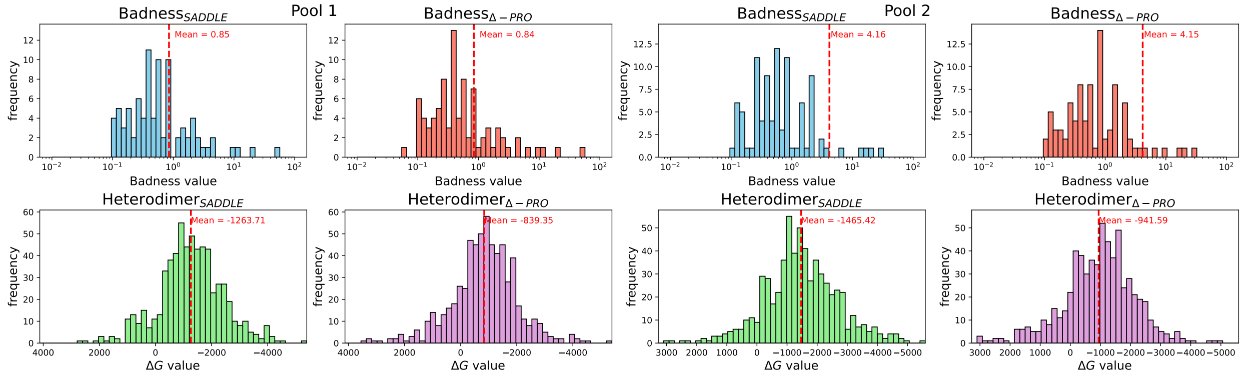


Figure 4: Results for primer candidates generated by Olivar (random seed 2340505846) on the FMD dataset.

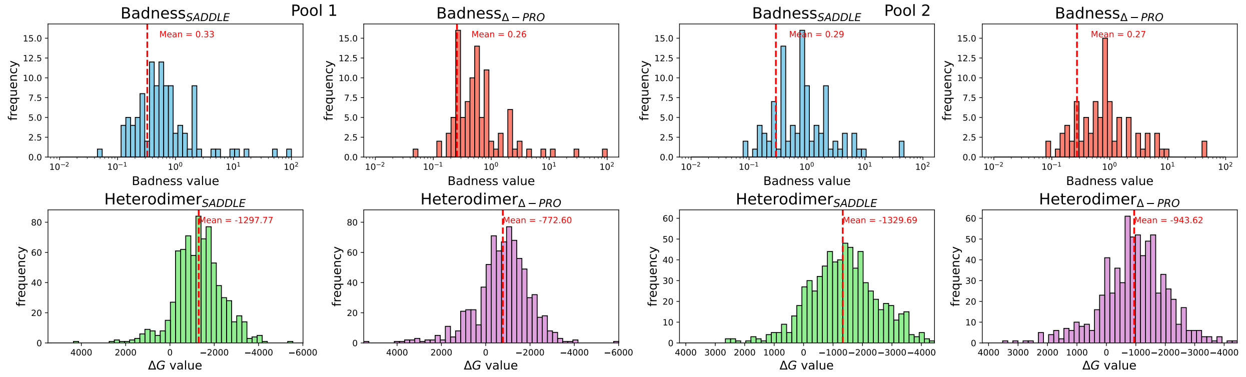


Figure 5: Results for primer candidates generated by Olivar (random seed 2530876844) on the FMD dataset.

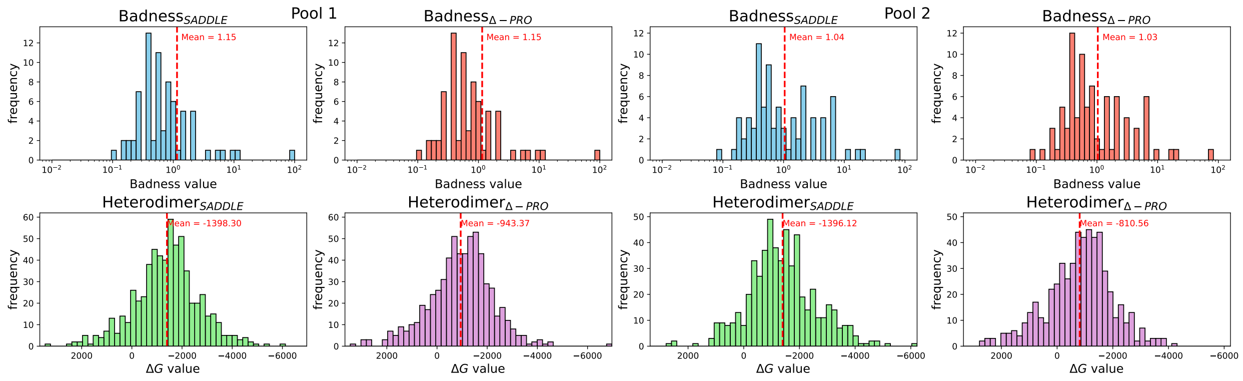


Figure 6: Results for primer candidates generated by Olivar (random seed 2746317213) on the FMD dataset.

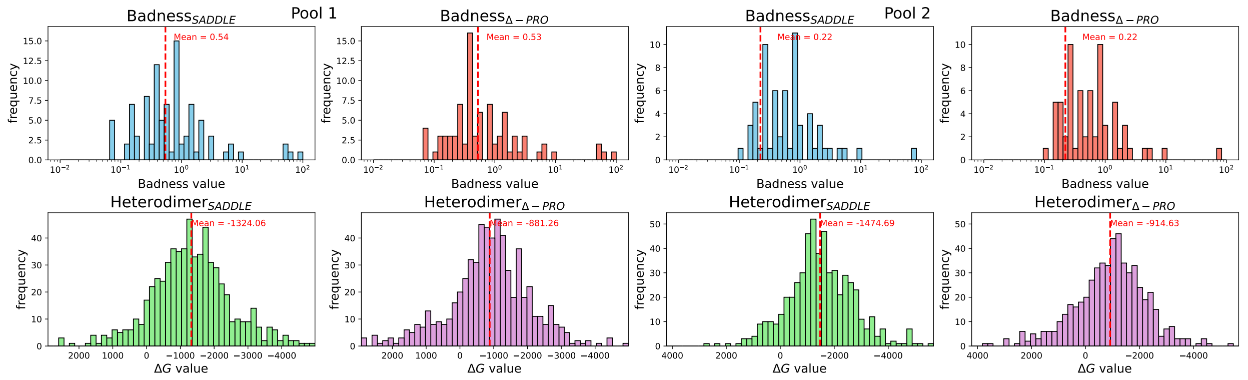


Figure 7: Results for primer candidates generated by Olivar (random seed 3163119785) on the FMD dataset.

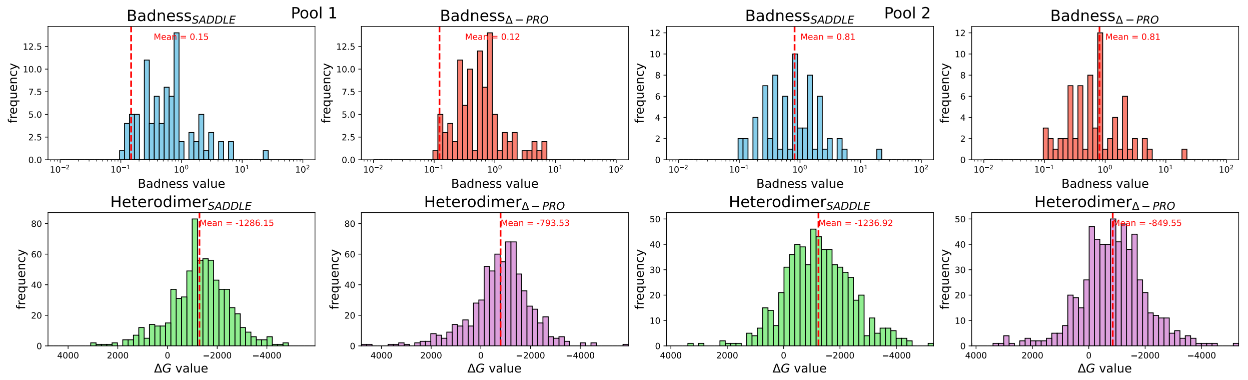


Figure 8: Results for primer candidates generated by Olivar (random seed 939042955) on the FMD dataset.

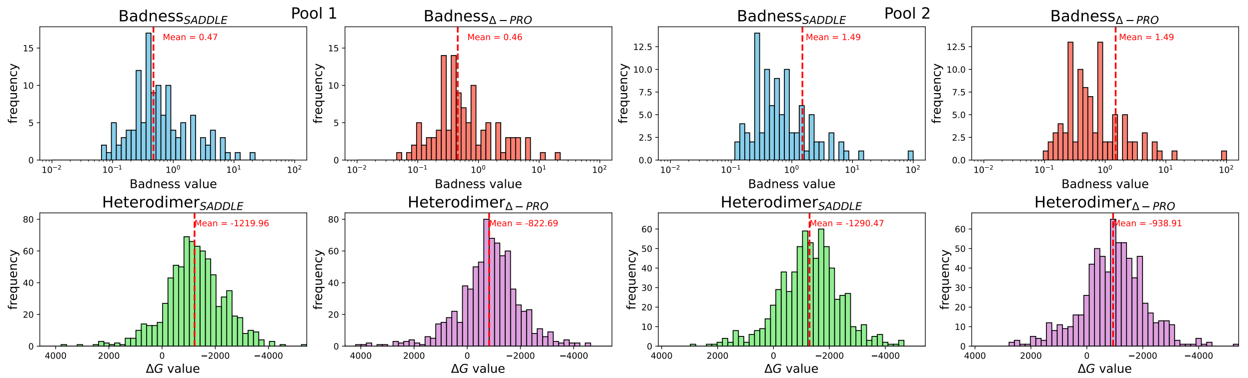

Figure 9: Results for primer candidates generated by Olivar (random seed 946785248) on the FMD dataset.

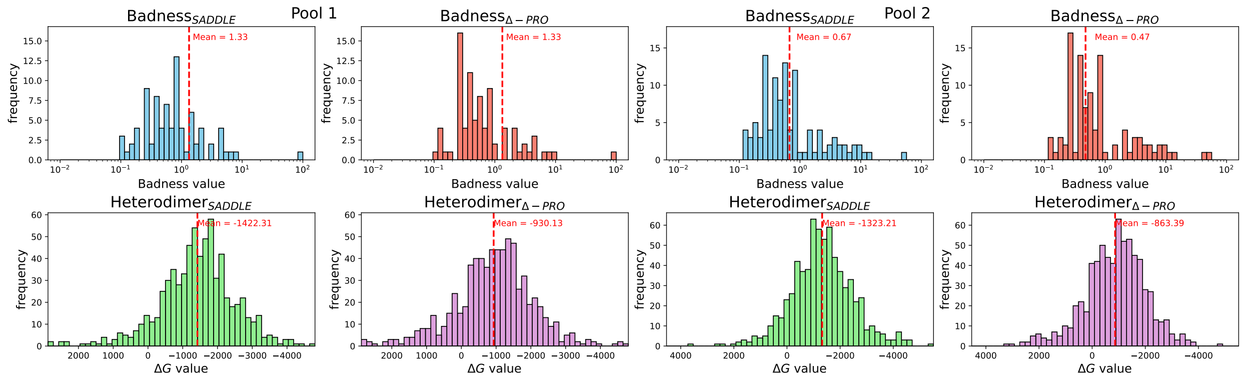

Figure 10: Results for primer candidates generated by Olivar (random seed 958682846) on the FMD dataset.

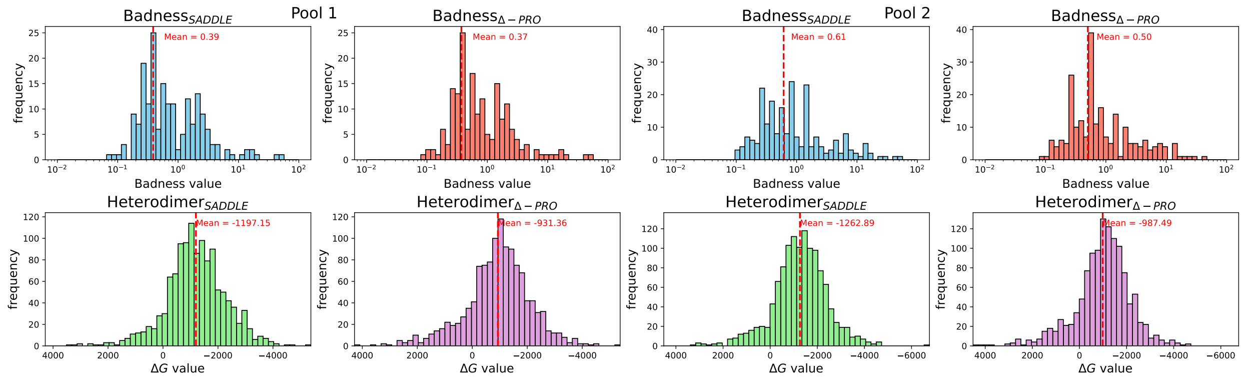

Figure 11: Results for primer candidates generated by Olivar (random seed 1181241943) on the ZIKA dataset.

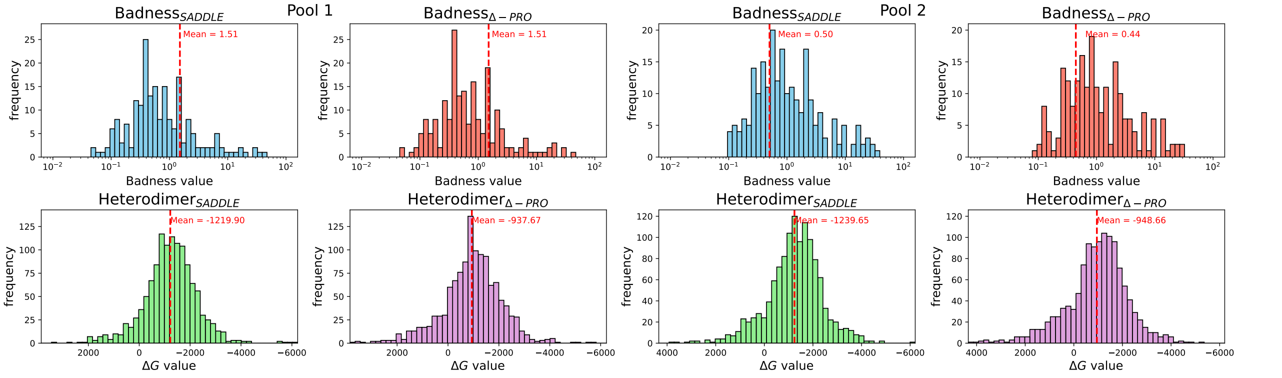


Figure 12: Results for primer candidates generated by Olivar (random seed 127978094) on the ZIKA dataset.

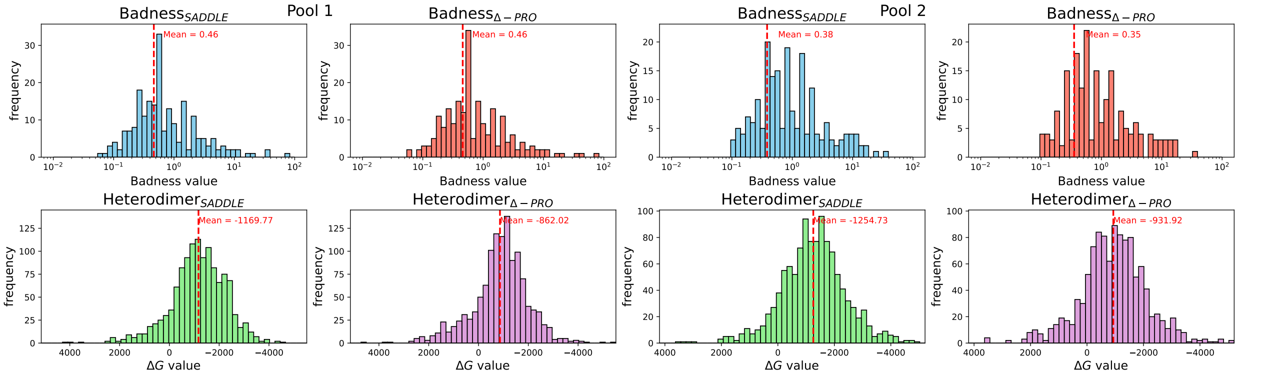


Figure 13: Results for primer candidates generated by Olivar (random seed 1812140441) on the ZIKA dataset.

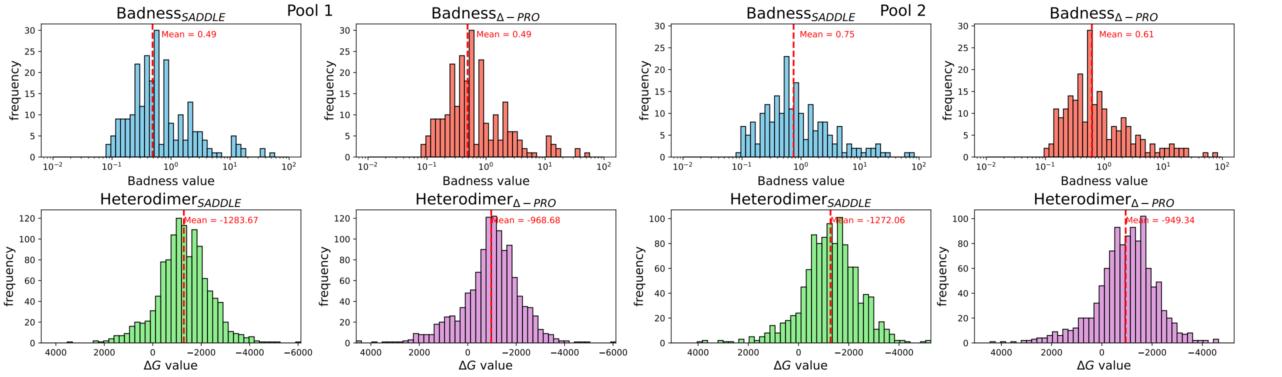


Figure 14: Results for primer candidates generated by Olivar (random seed 2340505846) on the ZIKA dataset.

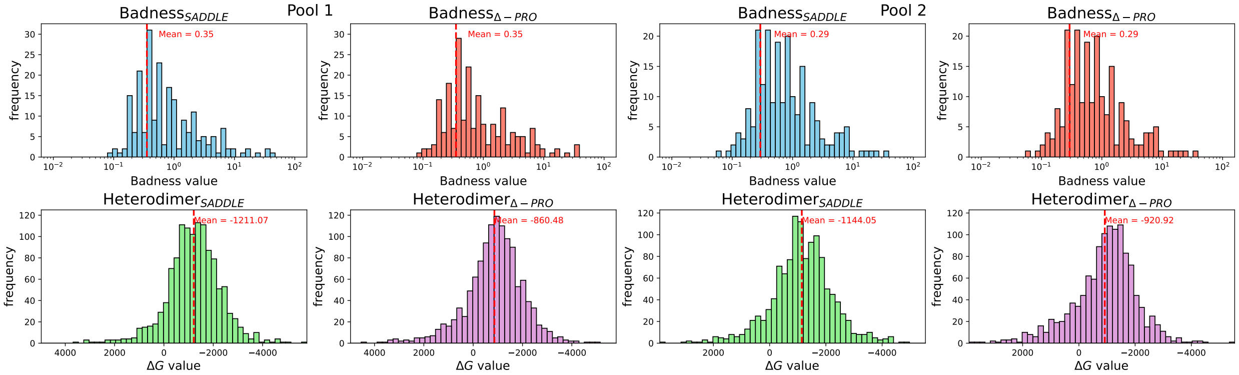


Figure 15: Results for primer candidates generated by Olivar (random seed 2530876844) on the ZIKA dataset.

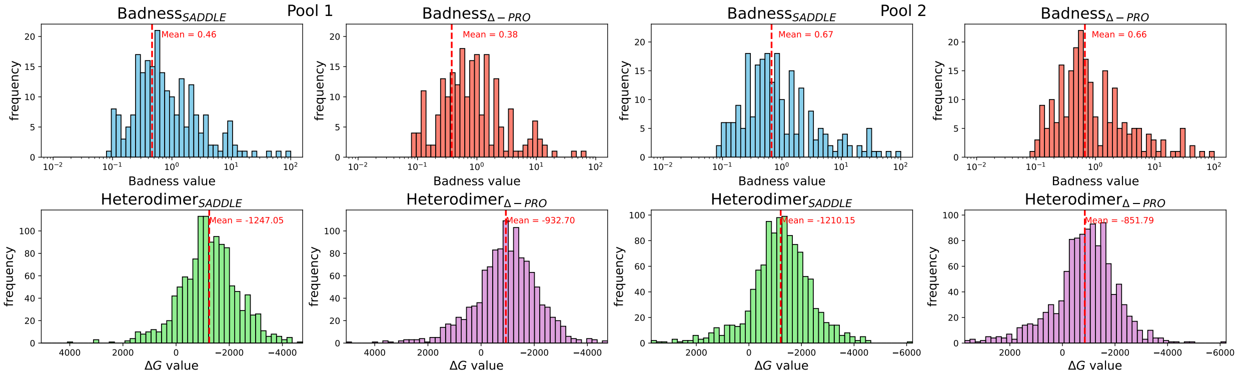


Figure 16: Results for primer candidates generated by Olivar (random seed 2746317213) on the ZIKA dataset.

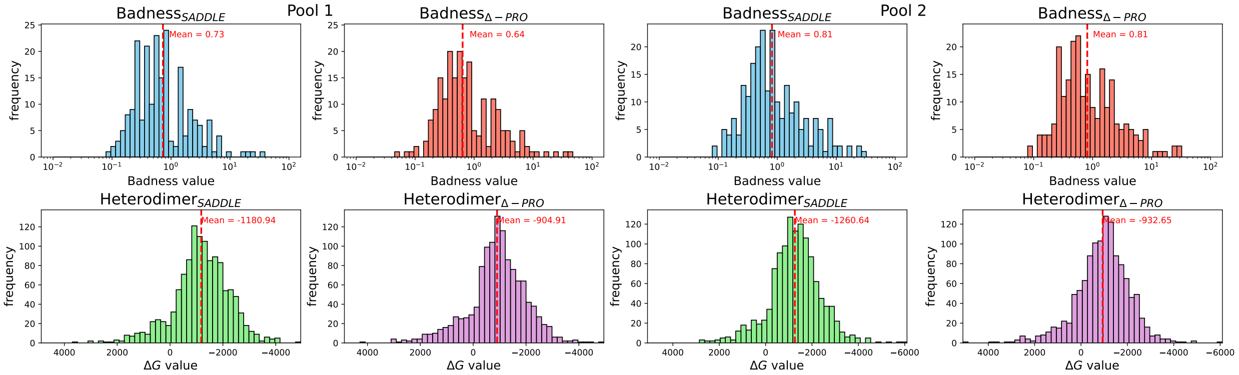


Figure 17: Results for primer candidates generated by Olivar (random seed 3163119785) on the ZIKA dataset.

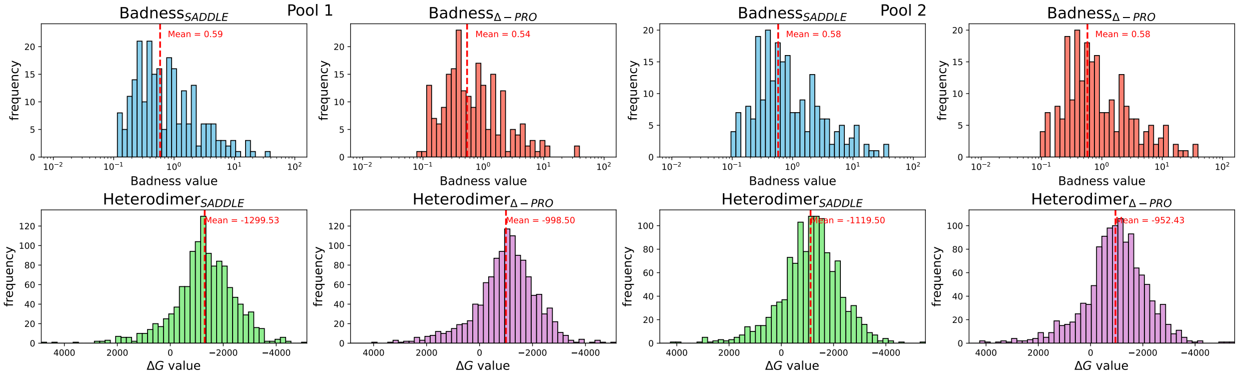


Figure 18: Results for primer candidates generated by Olivar (random seed 939042955) on the ZIKA dataset.

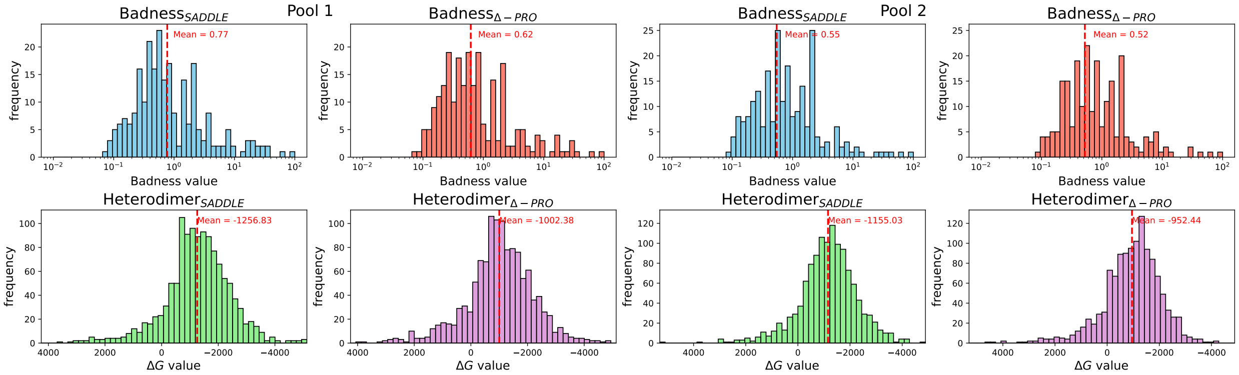


Figure 19: Results for primer candidates generated by Olivar (random seed 946785248) on the ZIKA dataset.

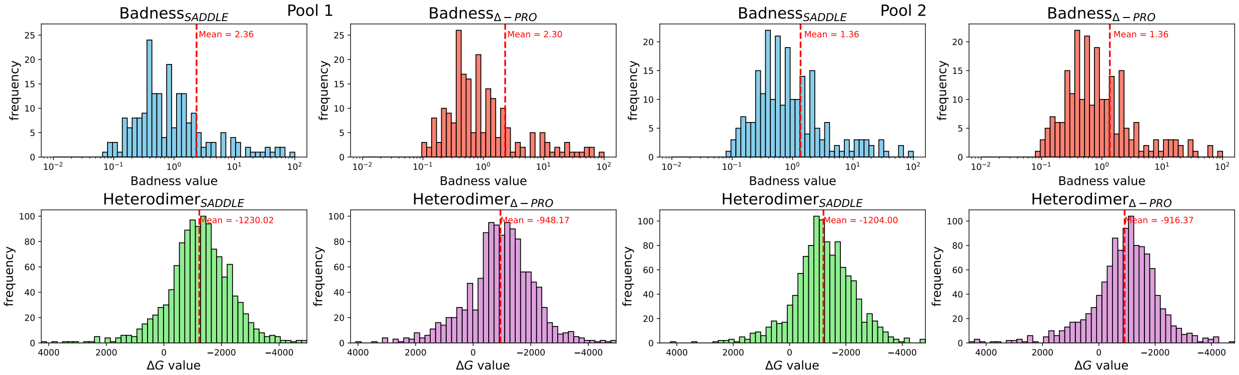


Figure 20: Results for primer candidates generated by Olivar (random seed 958682846) on the ZIKA dataset.
